# Adrenergic C1 neurons are part of the circuitry that recruits active expiration in response to hypoxia

**DOI:** 10.1101/774570

**Authors:** Milene R. Malheiros-Lima, Josiane N. Silva, Felipe C. Souza, Ana C. Takakura, Thiago S. Moreira

## Abstract

Breathing results from the interaction of two distinct oscillators: the preBötzinger Complex (preBötC) driving inspiration and the lateral parafacial region (pFRG) driving active expiration. The pFRG is silent during resting and become rhythmically active during high metabolic demand such as hypoxia. Catecholaminergic C1 cells are activated by hypoxia, which is a strong stimulus for active expiration. We hypothesized that the C1 cells and pFRG may constitute functionally distinct but interacting populations in order to contributes to control expiratory activity during hypoxia. We found that: a) C1 neurons are activated by hypoxia and project to pFRG region; b) active expiration elicited by hypoxia was blunted after blockade of ionotropic glutamatergic antagonist at the level of pFRG and c) selective depletion of C1 neurons eliminated the active expiration elicited by hypoxia. The results suggest that C1 cells may regulate the respiratory cycle including the active expiration under hypoxic condition.

## Introduction

The rostral ventrolateral medulla (RVLM) contains many types of neurons that regulate the sympathetic and parasympathetic outflows as well as breathing output (Guyenet, 2006; Del Negro *et al.*, 2018). Within the RVLM, respiratory physiologists named at least four functional segments, not counting the parafacial respiratory group/retrotrapezoid nucleus which can be viewed as the rostral-most extension of the so-called vental respiratory column (VRC) (Guyenet & Bayliss, 2015; Del Negro *et al.*, 2018); while cardiovascular physiologists named two crucial regions into the RVLM involved in sympathetic control. In the so-called "cardiovascular/sympathetic" area of the RVLM, the catecholaminergic C1 neurons was defined more than 40 years ago (for reviews (McAllen & Dampney, 1990; Guyenet, 2006; Guyenet *et al.*, 2013). C1 neurons have distinct projections to the whole brain, demonstrating that these neurons are involved in more than just cardiovascular regulation, as previously described (Ross *et al.*, 1981; Li & Guyenet, 1996; Schreihofer & Guyenet, 1997; Ritter *et al.*, 1998; Marina *et al.*, 2011; Abbott *et al.*, 2013*b*, 2013*a*, 2014). We and others have showed that selective stimulation of C1 cells causes a rise in arterial pressure and breathing activity (Abbott *et al.*, 2013*b*; Burke *et al.*, 2014; Malheiros-Lima *et al.*, 2018*a*). These effects caused by C1 stimulation mimic the cardiovascular and the respiratory responses elicited by hypoxia (Burke *et al.*, 2014). There is evidence that the C1 cells are highly collateralized, overlap with respiratory neurons in the VRC and contribute to hypoxic responses via connections with pontomedullary structures (Guyenet, 2006, 2014; Malheiros-Lima *et al.*, 2018*a*).

Hypoxia is considered very potent stimulus which produces different physiological perturbations including breathing activation (Guyenet, 2000; Prabhakar & Semenza, 2015). During hypoxic challenge, expiration (which is passive during exhalation) turns into an active process, with dilatation of upper airways and the recruitment of abdominal expiratory muscles during the second phase of the expiratory process (Iscoe, 1998). A group of expiratory neurons, located in the parafacial respiratory group (pFRG) are suggested to play a major role in the generation of active expiration (Janczewski & Feldman, 2006; Pagliardini *et al.*, 2011). The pFRG is located ventrolateral to the facial motor nucleus and becomes rhythmically active during periods of high metabolic demand such as hypoxia or hypercapnia (Pagliardini *et al.*, 2011; Huckstepp *et al.*, 2015).

Therefore, we postulate that the C1 neurons once activated recruit, via glutamatergic signaling, inspiratory (Malheiros-Lima *et al.*, 2018*a*) and expiratory (present results) muscles in an orderly sequence that is linked on the degree to which they are activated by hypoxia. This notion stems based on the following three observations: a) anatomically verified C1 neurons are activated by hypoxia and project to pFRG region; b) active expiration elicited by hypoxia was blunted after blockade of ionotropic glutamatergic antagonist at the level of pFRG and c) selective depletion of C1 neurons eliminated the active expiration elicited by hypoxia.

## Material and methods

### Animals

Experiments were performed in 35 adult male Wistar rats weighing 250-330 g. The animals were housed individually in cages in a room with controlled temperature (24 ± 2°C) and humidity (55 ± 10%). Lights were on from 7:00 am to 7:00 pm. Standard Bio Base rat chow (Águas Frias, SC, Brazil) and tap water were available *ad libitum*. Animals were used in accordance with the guidelines approved by the Animal Experimentation Ethics Committee of the Institute of Biomedical Science at the University of São Paulo (protocol number: 07/2014).

### Viral tracer injection into RVLM

We used a previously described lentiviral vector that expresses enhanced humanized channelrhodopsin-2 fused with eGFP under the control of the artificial Phox2b specific promoter PRSx8 (pLenti-PRSx8-hChR2 (H134)-eYFP-WPRE, henceforth abbreviated PRSx8-ChR2-eYFP) as a anterograde tracer (N = 4) (Hwang *et al.*, 2001; Abbott *et al.*, 2009*b*; Malheiros-Lima *et al.*, 2018*a*). Phox2b is a transcription factor expressed in subsets of brainstem neurons including the RTN and catecholaminergic C1 cells (Pattyn *et al.*, 1997; Stornetta *et al.*, 2006). The virus was produced by the virus core located at the Institute of Cancer under the supervision of Dr. Bryan Strauss (University of São Paulo) and diluted to a final titer of 6.75×10^11^viral particles/ml with sterile PBS before injection into brain. We verified that the above viral dilution produced specific transfection of catecholaminergic-expressing neuron. Under intraperitoneal (i.p.) injection of a mixture of ketamine (100 mg/kg) and xylazine (7 mg/kg) anesthesia, a glass pipette with external tip diameter 20 μm was inserted into the brain through a dorsal craniotomy. The lentivirus was delivered through the pipette by controlled pressure injection (60 PSI, 3-8 ms pulses). Stereotaxic coordinates to targeting C1 neurons were −2.8 mm caudal from lambda, ± 1.8 mm lateral from midline, and −8.4 mm ventral from the skull surface. The injection was targeted unilaterally to RVLM in a volume of 150 nL. (Malheiros-Lima *et al.*, 2018*a*). Animals were maintained for no less than 4 weeks before they were used in anatomical experiments. The surgical procedures, virus injections and guide cannula placement produced no observable behavioral or respiratory effects and these rats gained weight normally.

### Injection of retrograde tracer into parafacial respiratory group

In 5 animals, pFRG injections of the retrograde tracer, Cholera Toxin b (CTb, 1% in deionized water; List Biological, Campbell, CA) were unilaterally performed. For pFRG injections, the dorsal surface of the brain was exposed via an occipital craniotomy. Stereotaxic coordinates to target the pFRG were: −2.7 mm caudal from lambda; ±2.6 mm lateral from midline; −8.7 mm ventral from the skull surface. The CTb (30 nl) was delivered by pressure through glass pipettes. Retrograde tracers were injected over 1 min, and the pipette remained in the tissue for at least 5 min to minimize movement of tracer up the injection tract. The pipette was then removed, and the incision site closed. The animals were allowed 7-10 days for surgical recovery and for transport of the retrograde tracer.

### Immunotoxin lesions

Surgical procedures were performed on rats anesthetized with an i.p injection of a mixture of ketamine and xylazine (100 and 7 mg/kg of body weight, respectively). Postsurgical protection against infection included intramuscular injections of an antibiotic (benzylpenicillin, 160,000 U/kg). For selective chemical lesions of C1 cells, the rats were fixed to a stereotaxic frame and the coordinates used to locate the RVLM (−2.8 mm caudal from lambda; ±1.8 mm lateral from midline; −8.4 mm ventral from the skull surface) were based on the stereotaxic atlas for rats (Paxinos and Watson, 2007). The tip of a pipette, connected to a Hamilton syringe, was inserted directly into the RVLM for bilateral injections of saporin conjugate-dopamine beta hydroxylase (Anti-DβH-SAP; Advanced Targeting Systems, San Diego, CA) (2.4 ng in 100 nL of saline per side; N = 6). Based on a previous publication from our laboratory and the present study, we did not notice any differences in neuroanatomical or physiological experiments in animals that received IgG-saporin or saline in the C1 region (Taxini *et al.*, 2011; Malheiros-Lima *et al.*, 2017). Thus, in the present study, the sham-operated rats were injected with saline (0.15 M; N = 7). After surgery, the animals were kept in recovery for 2 weeks before they were used in physiological experiments.

### Physiological preparation

A tracheostomy was performed under general anesthesia with 5% isoflurane in 100% oxygen. Artificial ventilation (1ml/100g, 60-80 min^−1^) was initiated with 3.0-3.5% isoflurane in pure oxygen and maintained throughout surgical procedures. Frequency of ventilation was adjusted as needed to maintain end-tidal CO_2_ at the desired level. Arterial PCO_2_ was estimated from measurements of end-tidal CO_2_ and rectal temperature was maintained at 37±0.5°C (Guyenet *et al.*, 2005). This variable was monitored with a micro-capnometer (Columbus Instruments). Positive end-expiratory pressure (1.5 cm H_2_O) was maintained throughout to minimize atelectasis. Mean arterial pressure remained above 110 mmHg and the diaphragm activity could be silenced by lowering end-tidal PCO_2_ to 3.5-4.5%. In all animals, the femoral artery and vein were cannulated and both vagus nerves were cut distal to the carotid bifurcation as previously described (Malheiros-Lima *et al.*, 2018*a*). The diaphragm (Dia_EMG_) and abdominal (Abd_EMG_) muscles were accessed by a ventral approach, and muscles activities were performed before and after pharmacological manipulations within the pFRG region and chemoreflex challenges. Potassium cyanide (KCN: 40 μg/0.1 ml, i.v.) and 10% of the CO_2_ (balanced with O_2_) were used to activated chemoreceptors.

Upon completion of the surgical procedures, isoflurane was replaced by urethane (1.2 g/kg; iv.) administered slowly. The adequacy of anesthesia was monitored during a 20 min stabilization period by testing for the absence of withdrawal responses, pressor responses, and changes in Dia_EMG_ to a firm toe pinch. Approximately hourly supplements of one-third of the initial dose of urethane were needed to satisfy these criteria throughout the recording period.

### Physiological variables

As previously described, mean arterial pressure (MAP), diaphragm muscle activity (Dia_EMG_), abdominal muscle activity (Abda_EMG_), and end-expiratory CO_2_ (etCO2) were digitized with a micro1401 (Cambridge Electronic Design), stored on a computer, and processed off-line with version 6 of Spike 2 software (Cambridge Electronic Design, Cambridge, UK). Integrated diaphragm (∫DiaEMG) and abdominal (∫AbdEMG) muscle activity were obtained after rectification and smoothing (τ = 0.015s) of the original signal, which was acquired with a 30-300 Hz bandpass filter. Dia_EMG_ amplitude and frequency, Abd_EMG_ amplitude and MAP were evaluated before and after pharmacological manipulations.

### Drugs

All drugs were purchased from Sigma Aldrich (Sigma Chemicals Co.). Glutamate (10 mM - 50 nL; in sterile saline pH 7.4; unilateral) and kynurenic acid, a non-selective ionotropic glutamatergic antagonist (100 mM - 50 nL first dissolved in 1N NaOH and then diluted in phosphate-buffered saline pH 7.4; bilateral) were pressure injected (Picospritzer III, Parker Hannifin) (50 nL in 3 s) through single-barrel glass pipettes (20-μm tip diameter) into pFRG (Malheiros-Lima et al., 2018). All drugs contained a 5% dilution of fluorescent latex microbeads (Lumafluor, New City, NY, USA) for later histological identification of the injection sites.

### Hypoxia Protocol

Fos-like immunoreactivity evoked by hypoxia was studied in conscious, unrestrained adult rats. To acclimate the rats to the hypoxia environment prior to experimentation, they were kept in a plexiglass chamber (5 L) that was flushed continuously with a mixture of 79% nitrogen (N_2_) and 21% oxygen (O_2_) at a rate of 1 L/min, to allow them to become acclimated to the environmental stimuli associated with the chamber and to minimize unspecific fos expression. Rats were first acclimated for 45 min in the chamber and then subjected to acute hypoxia (AH) 8% O_2_, balanced with N_2_ or normoxic control 21% O_2_ for 3 hours, as previously demonstrated (King *et al.*, 2013; Silva *et al.*, 2016*b*). At the end of the stimulus, the rats were immediately deeply anesthetized and transcardially perfused. All experiments were performed at room temperature (24-26°C).

### Histology

The rats were deeply anesthetized with pentobarbital (60 mg/kg, i.p.), then injected with heparin (500 units, intracardially) and finally perfused through the ascending aorta, first with 250 mL of phosphate-buffered saline (PBS, pH 7.4) and then with 500 mL of 4% phosphate-buffered paraformaldehyde (0.1 M, pH 7.4). The brains were extracted, cryoprotected by overnight immersion in a 20% sucrose solution in phosphate buffered saline at 4 °C, sectioned in the coronal plane at 40 μm a sliding microtome and stored in cryoprotectant solution (20% glycerol plus 30% ethylene glycol in 50 mM phosphate buffer, pH 7.4) at −20 °C until histological processing. All histochemical procedures were completed using free-floating sections according to previously described protocols (Malheiros-Lima *et al.*, 2017, 2018*b*).

For immunofluorescence experiments we used the following primary antibodies: a) neuronal nuclei (NeuN) (mouse anti-NeuN antibody - 1:5000; Millipore, USA); b) tyrosine hydroxylase (TH) (mouse anti-TH - 1:1000; Chemicon, Temecula, CA, USA); c) ChR2-eGFP (chicken anti-GFP - 1:2000; Sigma, St. Louis, MO, USA); d) cholera toxin b (CTb) (goat anti-CTb - 1:1000; Chemicon, Temecula, CA, USA); e) vesicular glutamate transporter 2 (VGlut2, Slc17a6) (guinea-pig anti-VGlut2 - 1:2000; Chemicon International, Temecula, CA, USA) and f) fos (rabbit anti-fos - 1:2000; Santa Cruz Biotechnology, CA, USA). All of the primary antibodies were diluted in phosphate-buffered saline containing 1% normal donkey serum (Jackson Immuno Research Laboratories) and 0.3% Triton X-100 and incubated for 24 h. Sections were subsequently rinsed in PBS and incubated for 2 h in a) murine blue donkey anti-mouse (1:500) for NeuN; b) Alexa488 or Cy3 goat anti-mouse (1:200) for TH; c) Alexa488 or Cy3 donkey anti-chicken (1:200) for eGFP; d) Alexa488 goat anti-mouse (1:200) for CTb; e) Alexa 488 or Cy3 goat anti-guinea pig (1:200; Molecular Probes, USA) for VGlut2 and f) Cy3 goat anti-rabbit (1:200; Molecular Probes) for fos immunostaining. All the second antibody were from Jackson Laboratories (West Grove, PA, USA) unless otherwise stated. The sections were mounted on gelatin-coated slides in rostrocaudal sequential order, dried, and covered with DPX (Sigma Aldrich, Milwaukee, WI, USA). Coverslip were affixed with nail polish

### Mapping

A one in six series of 40 µm transverse sections through the brainstem were examined for each experiment under bright field and epifluorescence using a Zeiss AxioImager A1 microscope (Carl Zeiss Microimaging, Thornwood, NY). Neurons immunoreactive for NeuN, TH, eGFP, VGlut2, CTb, and/or fos were plotted with the Stereo Investigator software (Micro Brightfield, Colchester, VT) utilizing a motor driven microscope stage and the Zeiss MRC camera, after methods previously described (Takakura et al., 2006). Only cell profiles that included a nucleus were counted and/or mapped except in the cases where collaterals were mapped. The Stereo Investigator files were exported into the Canvas drawing software (Version 9, ACD Systems, Inc.) for text labeling and final presentation. The neuroanatomical nomenclature is after Paxinos and Watson (2007). Photographs were taken with a Zeiss MRC camera (resolution 1388 × 1040 pixels) and the resulting TIFF files were imported into the Canvas software. Output levels were adjusted to include all information-containing pixels. Balance and contrast were adjusted to reflect true rendering as much as possible. No other “photo-retouching” was done. Figures were assembled and labeled within the Canvas software.

The total number of TH (TH-ir), CTb and/or fos in the rostral ventrolateral medulla (RVLM: between 11.60 and 12.80 mm caudal to bregma level), commissural nucleus of the solitary tract (cNTS: between −14.28 and 15.96 mm caudal to bregma level), and parafacial respiratory group (pFRG: between −10.64 and −11.60) were plotted as mean ± SEM (RVLM: 6 sections/animal; cNTS: 8 sections/animal; pFRG: 5 sections/animal). The profile counts of the animals that received bilateral microinjections of anti-DβH-SAP reflected the sum of both sides of the brainstem and were compared with the control rats. The neuroanatomical nomenclature employed during experimentation and in this manuscript was defined by Paxinos and Watson (2007). The profile counts of animals that receive PRSx8-ChR2-eYFP or CTb injections reflected only one side of the medulla and were compared with the control rats. Confocal images (Carl Zeiss, Jena, Germany) were used to evaluate the colocalization between axonal varicosities of eGFP and VGlut2 in the pFRG region. Terminal fields were mapped using a x63 oil-immersion objective by taking 0.3 μm z-stack images of both red and green fluorescence through the tissue where discernible eGFP-labeled fibers were sharply in focus. These stacks were usually between 5 and 10 μm in depth. Terminals were marked as positive only where both eGFP and VGlut2 immunofluorescent profiles were in focus in at least 2 consecutive z sections.

### Statistics

All statistics were performed using GraphPad Prism 6. Data normality was assessed using the Shapiro-Wilk test, and the normally distributed data were expressed as means ± SEM. Statistical significance was accessed by student t-teste or two-way ANOVA, with repeated measures as appropriate with or without repeated measures. When applicable, the Bonferroni post-hoc test was used. Significance level was p<0.05.

## Results

### Active expiration elicited by activation of parafacial respiratory group is mediated by ionotropic glutamatergic receptors

Under resting condition, expiration occur passively, a consequence of the passive deflation of the lung and chest wall to resting from stretched positions at the end of inspiration (Figs. 1D-E and 1H). Physiologically, active expiration (high metabolic demand) corresponds to an increase in the airflow in the late expiratory phase immediately before inspiration, due to a recruitment of expiratory muscles. According to others, active expiration is present when Abd_EMG_ amplitude during the final 20% of the expiratory period was >50% larger than in the first half (Figs. 1D-E) (Pagliardini *et al.*, 2011; Zoccal *et al.*, 2018; Silva *et al.*, 2019). In adult rat, the pFRG region (a conditional expiratory oscillator) is inactive at rest because of the intense synaptic inhibition (Janczewski & Feldman, 2006; Pagliardini *et al.*, 2011).

**Figure 1).**
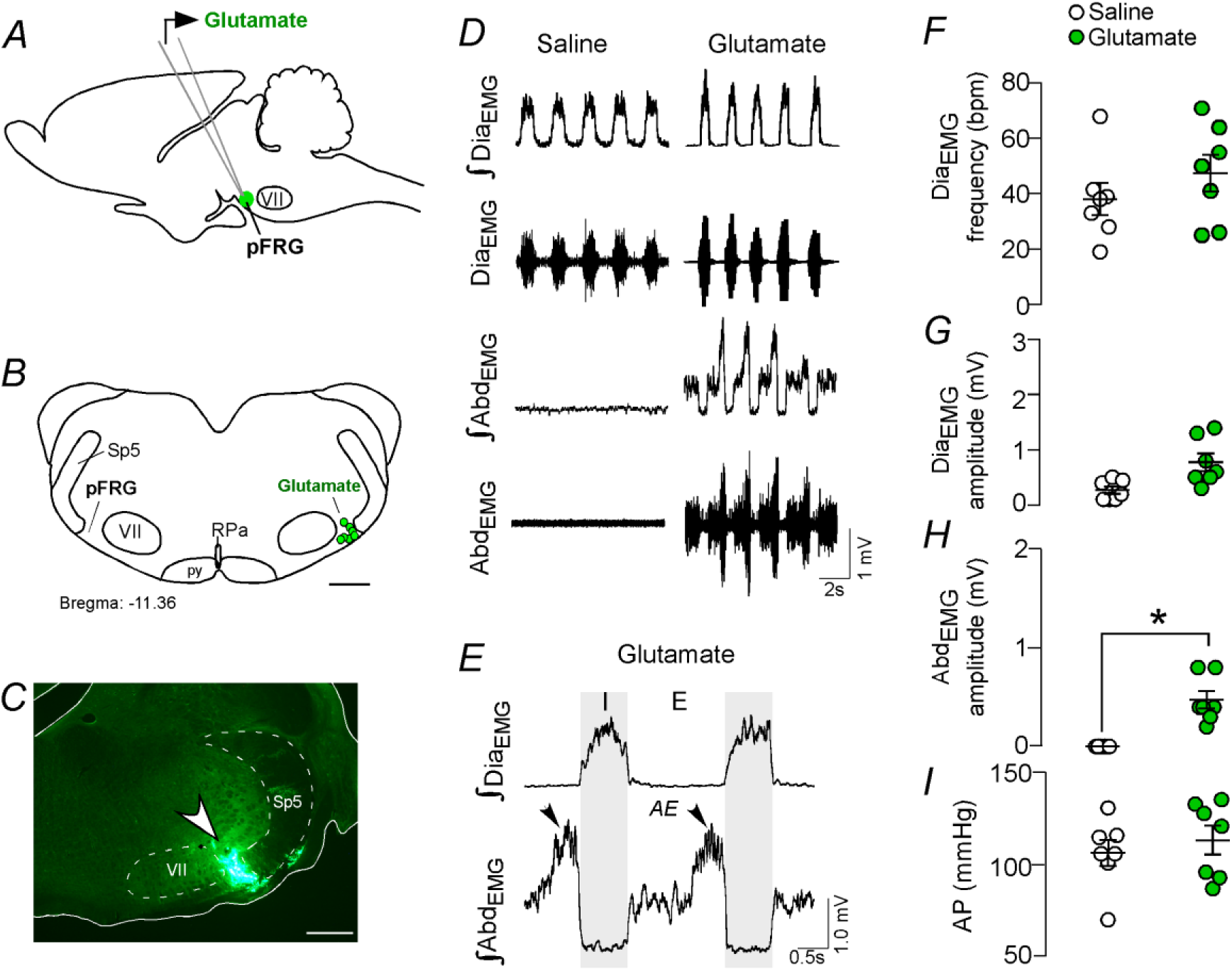
Activation of glutamatergic receptors into pFRG region evoked active expiration. A) Experimental design. B) Injection sites of glutamate (10 mM - 50 nL) into the pFRG region. C) Photomicrography showing a typical injection site of glutamate into pFRG region. D) Traces showing breathing parameters from a representative experiment. E) Expanded traces after glutamate injection into the pFRG. Black arrows show active expiration (AE) in the expanded traces. F-I) Individual data for each parameter: F) Dia_EMG_ frequency (bpm), G) Dia_EMG_ amplitude (mV), H) Abd_EMG_ amplitude (mV) and I) AP (mmHg) after unilateral injections of saline (white circles) or glutamate (green circles) into pFRG. Lines show mean and SEM. *Different from saline, Paired Student’s t-test, p<0.05. Abbreviations: pFRG, parafacial respiratory group; Sp5, spinal trigeminal tract; VII, facial motor nucleus; Dia_EMG_, diaphragm electromyogram; Abd_EMG_, abdominal electromyogram; bpm, breaths per minute; AP, arterial pressure; AE, active expiration. Scale bar in B = 1mm and C = 0.5mm.

Considering the idea that pFRG neurons are involved in active expiration and C1 neuron use glutamate as main neurotransmitter, the first question that we address was: Does activation of ionotropic glutamatergic receptors in the pFRG neurons able to generate active expiration?

We performed unilateral injections of glutamate (10 mM - 50 nL; N = 7) (Figs. 1A-C), followed by ipsilateral injections the broad spectrum ionotropic glutamatergic antagonist kynurenic acid (kyn, 100 mM - 50 nL; N = 7) into the pFRG region. According to our histological analysis, the injections were located in the ventral aspect of the lateral edge of the facial nucleus, juxtaposed to the spinal trigeminal tract (Figs. 2B-C). Unilateral injection of glutamate into the pFRG triggered active expiration, as noted by bursts of Abd_EMG_ during late expiration (Figs. 1D-E). For example, glutamate into the pFRG elicited and increase in Abd_EMG_ (0.236±0.045, vs. baseline: 0.000±0.000 mV, p<0.05) (Figs. 1D-E and 1H). The injection of glutamate into the pFRG did not evoke an increase in Dia_EMG_ frequency (47±7, vs. baseline: 38±6 bpm, p>0.05), Dia_EMG_ amplitude (0.771±0.160, vs. baseline: 0.264±0.068 mV, p>0.05) nor blood pressure (124±7, vs. baseline: 120±7 mmHg, P>0.05) (Figs. 1D-I). Previous ipsilateral injection of kyn into the pFRG completely abolish the active expiration generated by glutamate injections (0.000±0.000, vs. glutamate: 0.236±0.045 mV, p<0.05). Blockade of the ionotropic glutamatergic receptors at the level of pFRG did not affect resting Dia_EMG_ frequency (38±5 vs. glutamate: 47±7 bpm, p>0.05), Dia_EMG_ amplitude (0.3142±0.114, vs. glutamate: 0.771±0.160mV, p>0.05) nor blood pressure (127±8, vs. glutamate: 124±7mmHg, p>0.05).

**Figure 2).**
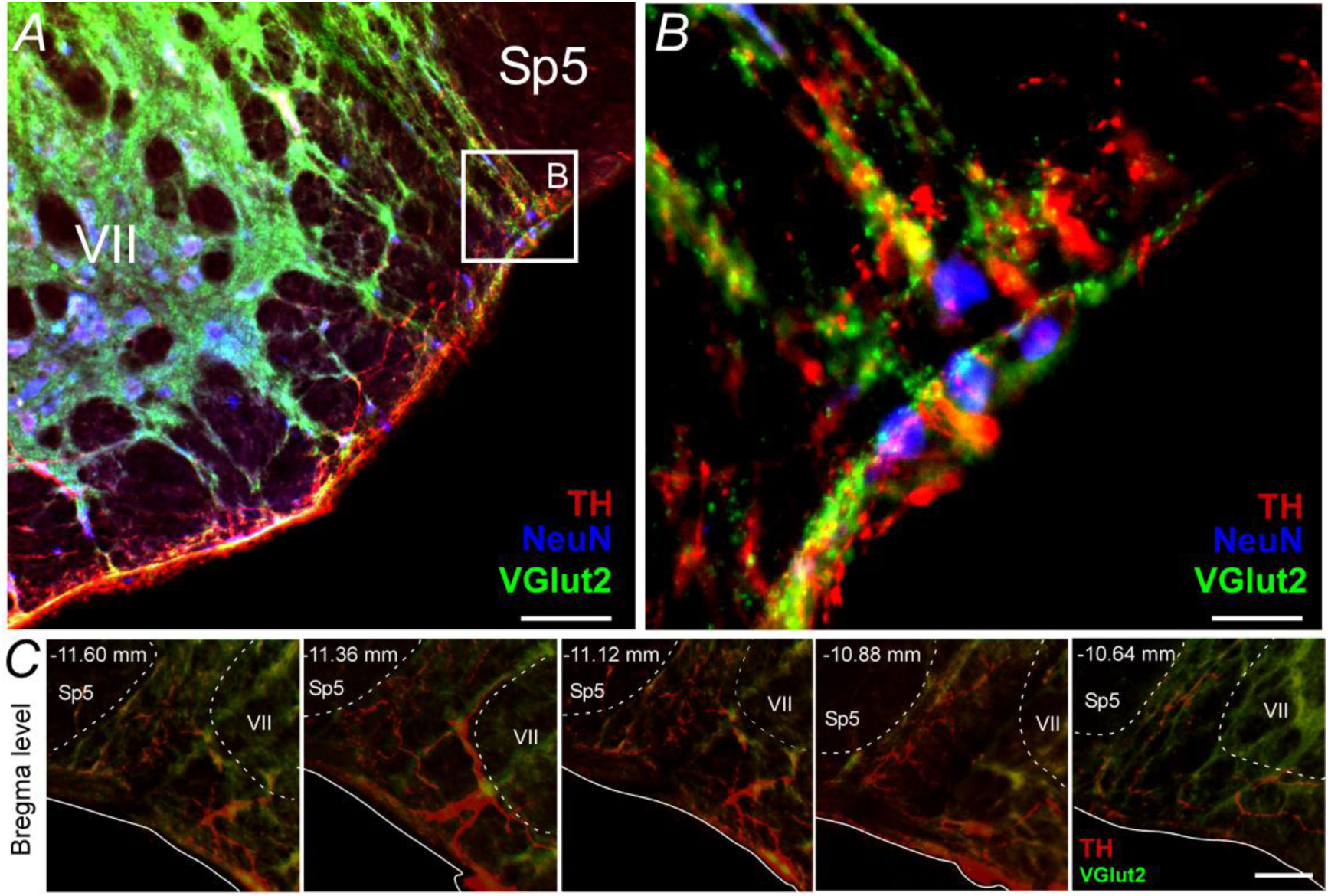
Glutamatergic and catecholaminergic innervations are located into the pFRG region. A-B) Single case showing the close apposition between glutamatergic (VGlut2; green) and/or catecholaminergic (TH; red) terminals in the pFRG neurons (NeuN; blue). C) Rostro-caudal distribution of glutamatergic and catecholaminergic terminal in the pFRG region (Bregma level: −11.60 at 10.64 mm). Note the overlap between VGlut2 and TH terminals. Abbreviations: Sp5, spinal trigeminal tract; VII, facial motor nucleus; TH, tyrosine hydroxylase; VGlut2, vesicular glutamate transporter 2; NeuN, nuclear neuronal marker. Scale bar in A= 1 mm, B = 10 μm, and C = 100 μm.

### Catecholaminergic and glutamatergic terminals at the level of the parafacial respiratory group

Based on the fact that activation of ionotropic glutamatergic receptors elicit active expiration (Fig. 1 and (Huckstepp *et al.*, 2018) and the existence of a glutamatergic and/or catecholaminergic mechanism for the generation of active expiration during hypoxia (Malheiros-Lima *et al.*, 2017, 2018*a*), our second series of experiments was to investigate the presence of catecholaminergic innervation in the region of the pFRG. Coronal brainstem sections were immunolabelled for tyrosine hydroxylase (TH) and vesicular glutamate transporter (VGlut2), two markers that respectively identify catecholaminergic and glutamatergic fibers and terminals in the pFRG region (Figs. 2A-C). Catecholaminergic and/or glutamatergic terminals and fibers were observed lateral to the caudal edge of the facial nucleus, a region described as a conditional expiratory oscillator (Fig. 2). Within the pFRG, we observed several neurons (NeuN^+^) in close contact with TH^+^ and/or VGlut2^+^ terminals located lateral to the caudal tip of the facial nucleus (Fig. 2B).

### Parafacial respiratory group targeted by ChR2-expressing catecholaminergic C1 neurons

In order to be more specific about the projection of catecholaminergic C1 neurons to the pFRG region, the next series of experiment was designed to identify the presence of axonal varicosities in the pFRG region. Injections of the lentivirus PRSx8-ChR2-eYFP into the left C1 region produced intense fluorescence protein expression only in neurons (Figs. 3A; 3C and 3C’). These neurons (TH^+^/eYFP^+^) were always located in close proximity to the original injection site and were concentrated in the region of the ventrolateral reticular formation that lies below the caudal end of the facial motor nucleus and extends up to 500 μm posterior to this level (−11.6 to - 12.8 mm caudal to Bregma; Figs. 3B-C). This region contains the bulk of the C1 and other blood-pressure regulating presympathetic neurons (Dampney, 1994; Guyenet, 2006; Guyenet *et al.*, 2013). Smaller injections were performed to avoid the transduction of the retrotrapezoid nucleus (RTN) and A1 region (data not shown; Malheiros-Lima *et al.*, 2018). Figure 3B show the total number of TH^+^ neurons expressing GFP for 4 rats used in the tracer experiments. On average, 56 ± 3% of the TH-positive neurons were immunoreactive (ir) for GFP (Fig. 3B).

**Figure 3).**
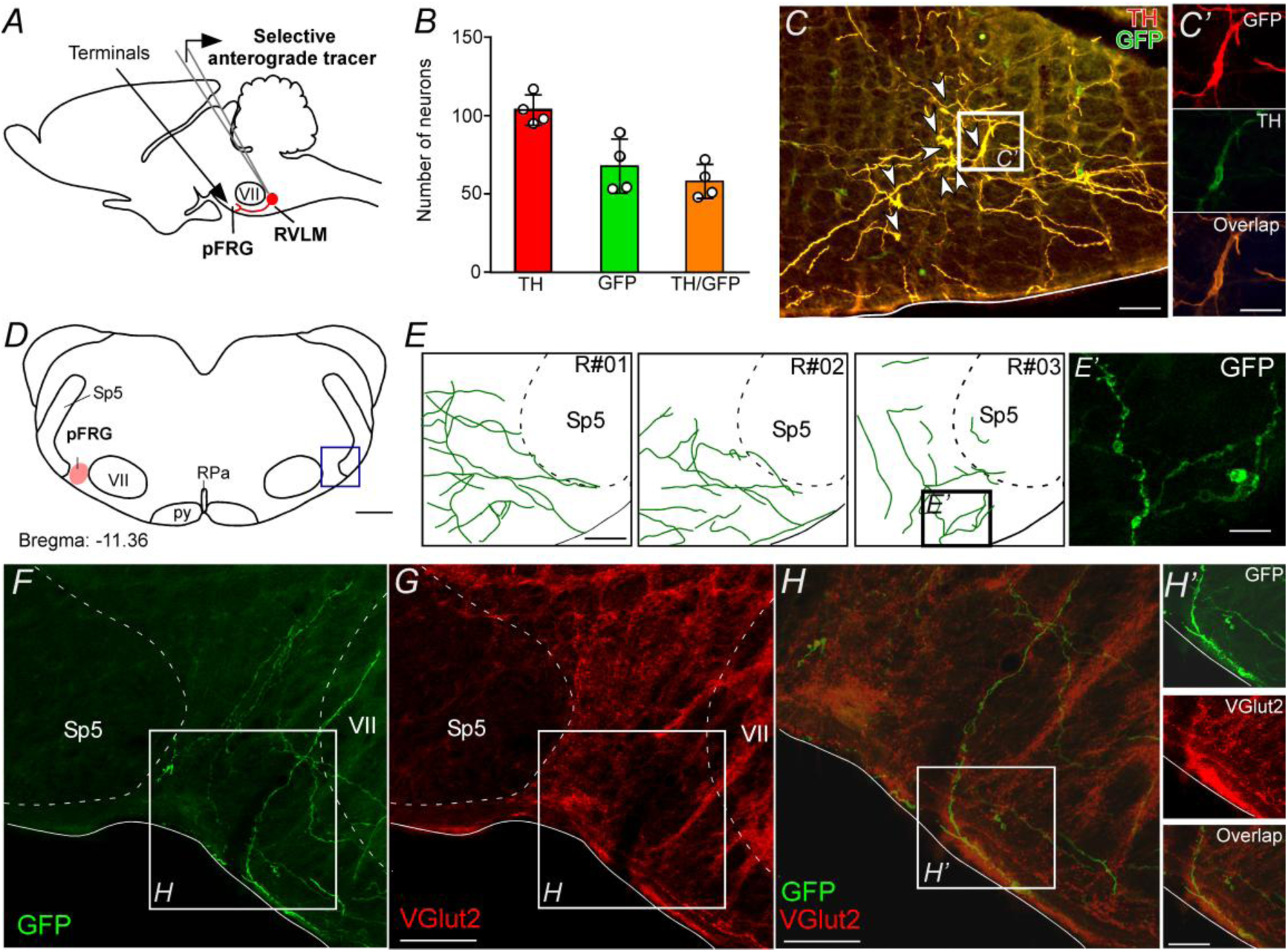
pFRG receives glutamatergic inputs from catecholaminergic C1 neurons. A) Experimental design. B) The number of TH and GFP neurons in the RVLM (N = 4). Cell count was obtained in 6 coronary brain sections (40 μm with 240 μm intervals between slices) from each rat. C-C’) Sigle case showing that the C1 neurons are transfected by the lentivirus PRSX8-ChR2-eYFP. D-E) Schematic drawing of a coronal brain section showing the distribution of GFP terminals in the pFRG region in 3 rats (bregma level: −11.36 mm according with Paxinos and Watson atlas, 2007). E’) Photomicrography showing terminals into the pFRG region. F-H’) Photomicrography showing fibers and terminals expressing lentivirus (GFP) and glutamate (VGlut2) in the pFRG region. Abbreviations: pFRG, parafacial respiratory group; RVLM, rostral ventrolateral medulla; Sp5, spinal trigeminal tract; VII, facial motor nucleus; FN, facial nerve; RPa, raphe pallidus; py, – pyramid tract; TH, tyrosine hydroxylase; GFP, green fluorescent protein; VGlut2, vesicular glutamate transporter 2. Scale bar in C and E-H = 50μm, C’, E’ and H’= 20μm, D = 1mm.

Transduced catecholaminergic neurons had putative catecholaminergic and glutamatergic varicosities within the pFRG (Figs. 2; 3D-H; 3E’ and 3H’). We also noticed that every pontomedullary region that harbors noradrenergic neurons including the locus coeruleus, A1, A2 and A5 region also have GFP labeled fibers (data not shown). We observed light projections within the medullary raphe, a region also noted for its role in respiratory control, moderately dense projections to the nucleus of the solitary tract which receives input from the carotid bodies and other cardiopulmonary afferents, and extremely dense projections to the dorsal motor nucleus of the vagus. All projections had a strong ipsilateral predominance.

### Parafacial respiratory group-projecting catecholaminergic neurons in the C1 region

Five cases with CTb injections within the pFRG were analyzed to visualize retrograde labeling in the C1 region (Figs. 4C and 4E). A Representative CTb injections centered on the pFRG of one rats is shown in figure 4B. C1 neurons with projections to pFRG were labeled with CTb seven to ten days before the rats were sacrificed. TH immunoreactivity was used to identify C1 neurons (Figs. 4C and 4E). To characterize the C1 neurons that project to pFRG, a series of 40 μm-thick coronal were analyzed and cells were counted in only 6 levels of the C1 region: −11.6 to - 12.80 mm relative to bregma, in order to include the lowest possible amount of catecholaminergic neurons in the A1 neurons. A considerably high number of retrograde labeled neurons (CTb^+^) were found in the C1 region (Figs. 4C and 4E), particularly in more rostral regions of the C1 region. For example, we found from a total of 129±11 TH-ir cells in the C1 region, 107±10 neurons project to pFRG, representing 83% of TH cells in the C1 region (Figs. 4C and 4E). As expected, CTb expression was also observed in cell bodies located in the commissural aspect of the nucleus of the solitary tract (cNTS); however few double labeled cells (49±4 vs. total TH: 148±9; 33% of TH cells in the A2 region) were found within the cNTS (Figs. 4D and 4F).

**Figure 4).**
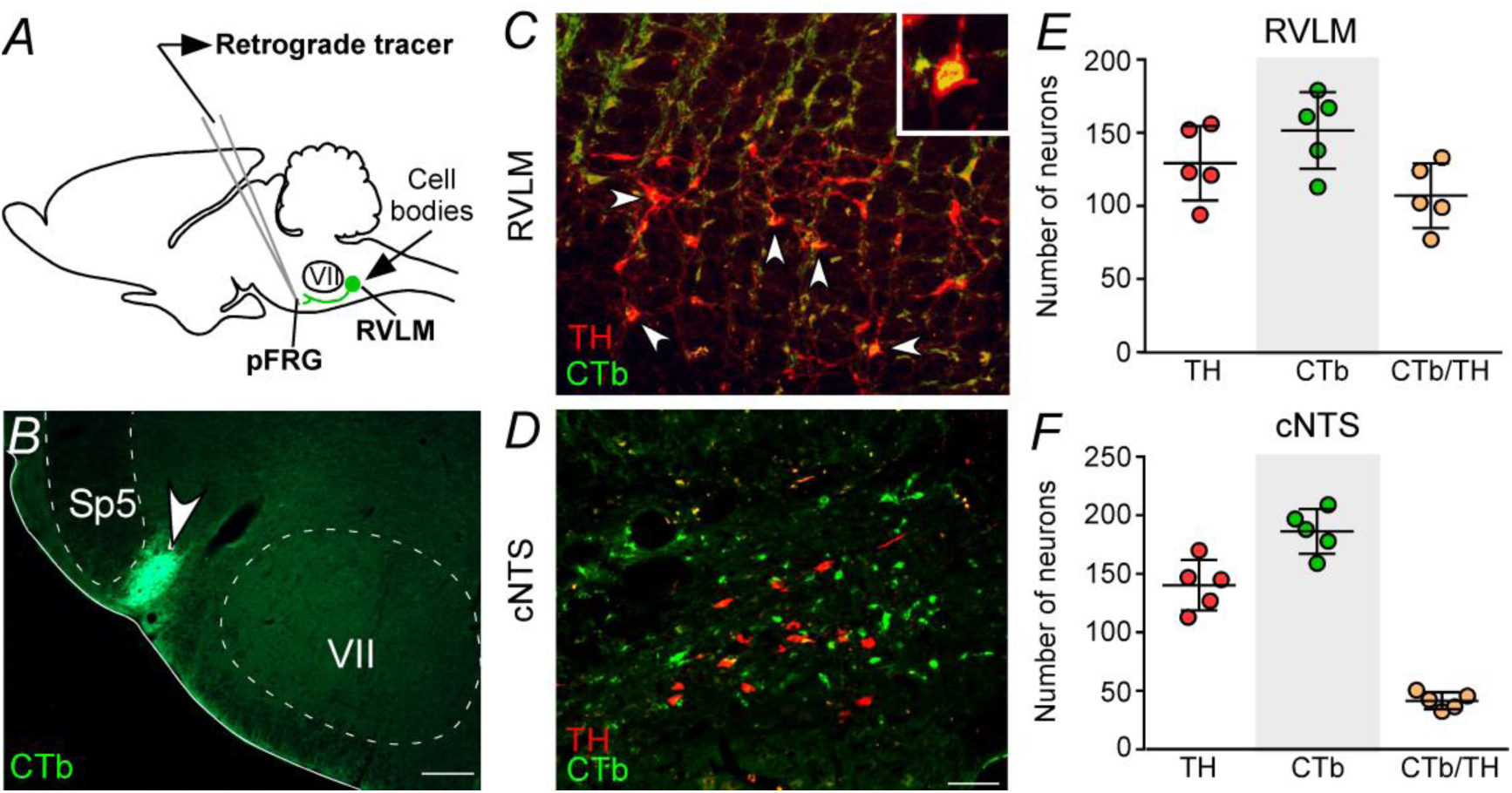
Catecholaminergic C1 neurons projects to the pFRG region. A) Experimental design. B) Single case showing a positive injection of the retrograde tracer CTb (1%, 30-50 nL) into the pFRG region. C-D) Photomicrography of catecholaminergic (TH) and CTb-labelled cells located in the RLVM and cNTS of one rat with CTb positive injection into pFRG. E-F) Number of CTb and/or TH-labeled cells in the RVLM and cNTS. Cell count was obtained in coronary brain sections (RVLM: 6 sections and cNTS 8 sections: 40 μm with 240 μm of intervals between slices) from each rat. Abbreviations: pFRG, parafacial respiratory group; Sp5, spinal trigeminal tract; VII, facial motor nucleus; RVLM, rostral ventrolateral medulla; CTb, Cholera Toxin b; TH, tyrosine hydroxylase; cNTS, commissural nucleus of the solitary tract. Scale bar in B = 0.5 mm and C-D = 50 μm.

### Parafacial respiratory neurons are activated by hypoxia

In order to verify if neurons located into the pFRG region are activated by hypoxia, fos expression was evaluated in rats exposed to hypoxia (8% O_2_, balanced with N_2_) or normoxia (21% O_2_, 78% N_2_, 1% CO_2_) for 3 hours. Coronal brainstem sections were double labelled for fos (used as a reporter of cell activation) and tyrosine hydroxylase (TH, marker for catecholaminergic cell bodies and terminals). The number of fos and TH neurons were identified and counted in a one-in-six series of transverse sections (1 section every 240 μm). Counts were made on both sides of the brain and throughout the portion of the rostral aspect of the ventrolateral medulla (pFRG and C1 regions) (Figs. 5A-D; 5G-H). We also counted the number of fos^+^ neurons located in the cNTS)(Figs. 5E-F; 5I).

**Figure 5).**
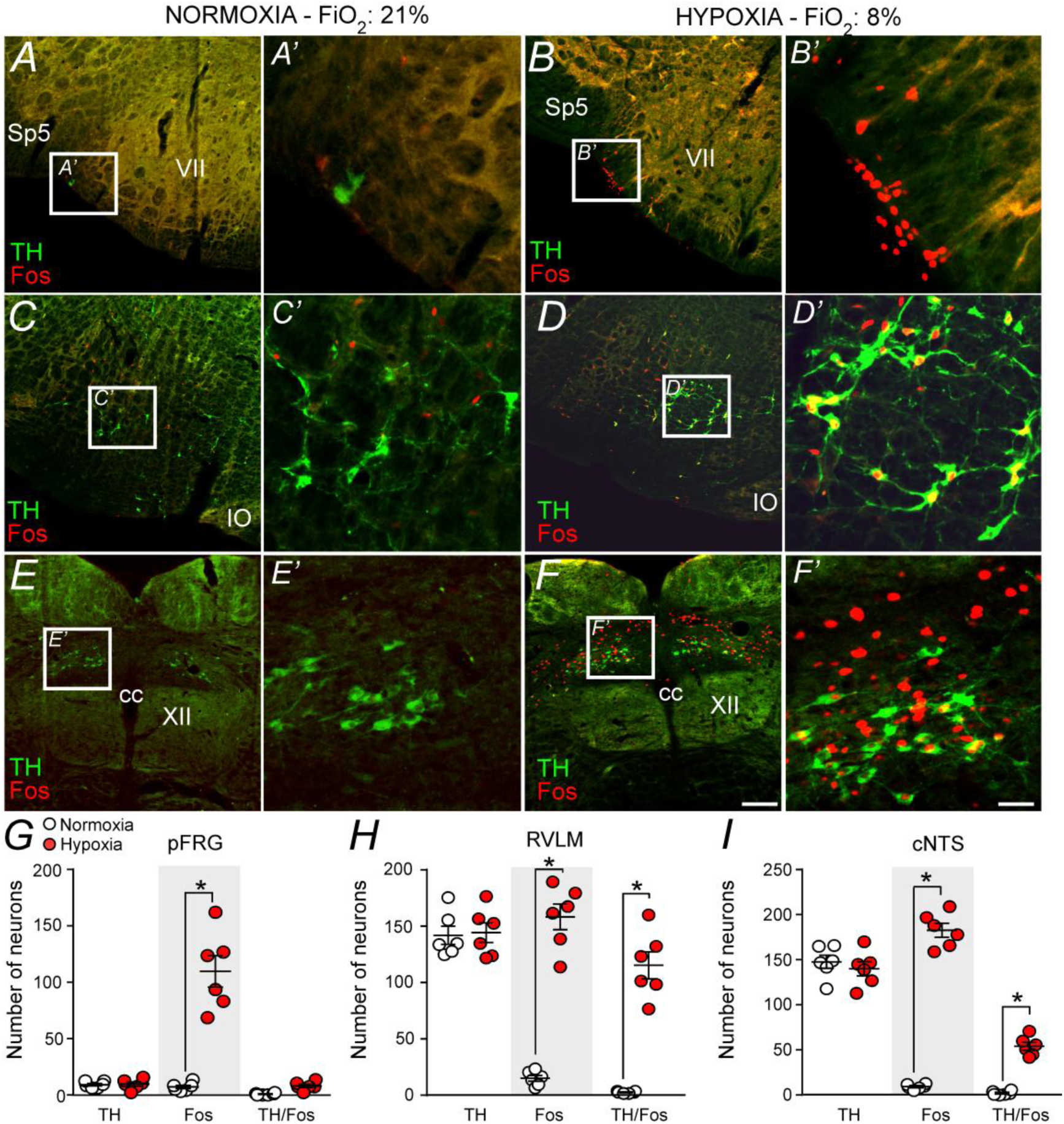
Hypoxia triggers neurons in the pFRG, RVLM and cNTS. A-F) Photomicrography showing the representative cases of the TH and Fos-labelled cells located in the (A) pFRG, (B) RLVM and (C) cNTS after normoxia (21% O_2_, balanced with N_2_) and after hypoxia (3 hours in 8% O_2_, balanced with N_2_). A’-F’) Higher magnification of pFRG, RVLM and cNTS showing neurons labelling to Fos and/or TH. G-I) Number of Fos and/or TH-labeled cells in the pFRG, RVLM and cNTS. Cell count was obtained in coronary brain sections (pFRG: 6 sections; RVLM: 6 sections and cNTS 8 sections: 40 μm with 240 μm of intervals between slices) from each rat. Abbreviations: pFRG, parafacial respiratory group; RVLM, rostral ventrolateral medulla; cNTS, commissural nucleus of the solitary tract; Sp5, spinal trigeminal tract; VII, facial motor nucleus; IO, inferior olive; cc, central canal; XII, hypoglossal nucleus; TH, tyrosine hydroxylase. Scale bar in A-F = 100 μm and A’-F’ = 20μm.

Figure 5A shows the pFRG region in a control rat (normoxia) and Figure 5B in a rat exposed to hypoxia in a period of 3 hours. Hypoxia caused a large increase in the number of fos^+^ cells in presumably pFRG neurons (fos^+^/TH^−^) compared to normoxia (control group) (115±14, vs. normoxia: 7±2, p<0.001) (Figs. 5A-B, and G). As expected, we also detected a large amount of fos^+^ cells in C1 neurons (fos^+^/TH^+^) (158±11, vs. normoxia: 14±3, p<0.001) (Figs. 5C-D, and 5H). The number of fos^+^ neurons in the cNTS (183±8, vs. normoxia: 10±2, p<0.001) were higher in hypoxia group compared to normoxia group (Figs. 5E-F, and 5I).

### Effect of bilateral blockade of ionotropic glutamatergic receptors in the parafacial respiratory group on active expiration elicited by chemoreflex activation

The follow experiment was designed to determine whether the activation of pFRG was able to elicit active expiration during hypoxia or hypercapnia depends on glutamatergic synapses. Bilateral Kyn (100 mM - 50 nL) injections were then performed in the pFRG (Figs. 6A-C), and the effects on peripheral chemoreflex activation (bolus injection of KCN)-induced changes in Dia_EMG_, Abd_EMG_ and MAP were evaluated (Figs. 6D-I).

**Figure 6).**
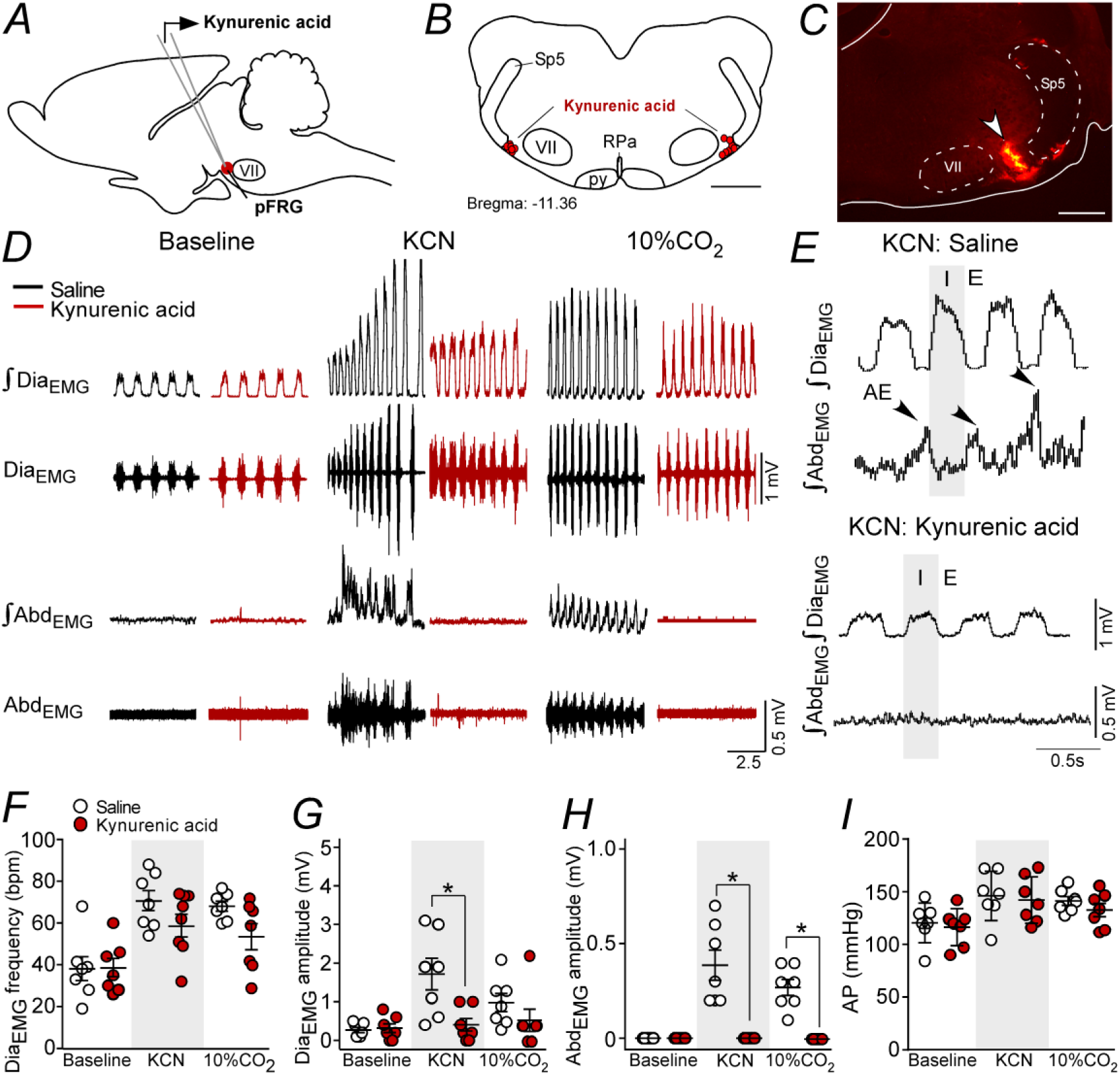
Kynurenic acid injection into pFRG region blunted active expiration induced by hypoxia and hypercapnia. A) Experimental design. B) Bilateral injection sites of kynurenic acid (100 mM - 50 nL) into pFRG. C) Photomicrography showing a typical injection site of kynurenic acid into pFRG. D) Traces showing breathing parameters from a representative experiment. E) Expanded traces. Black arrows show active expiration (AE) in the expanded traces. F-I) Individual data for each parameter: F) Dia_EMG_ frequency (bpm), G) Dia_EMG_ amplitude (mV), H) Abd_EMG_ amplitude (mV) and I) AP (mmHg) after bilateral injections of saline (white circles) or kynurenic acid (red circles) into pFRG. Lines show mean and SEM. *Different from saline; Two-way ANOVA, p<0.05. Abbreviations: pFRG, parafacial respiratory group; Sp5, spinal trigeminal tract; VII, facial motor nucleus; Dia_EMG_, diaphragm electromyogram; Abd_EMG_, abdominal electromyogram; bpm, breaths per minute, AP, arterial pressure; KCN, potassium cyanide. Scale bar in B = 1 mm and C = 0.5 mm.

As expected, the antagonism of glutamatergic receptors in the pFRG with Kyn injections did not alter resting Dia_EMG_ frequency (38±5, vs. baseline: 38±6 bpm, p>0.05), Dia_EMG_ amplitude (0.314±0.114, vs. baseline: 0.264±0.067 mV, p>0.05) nor blood pressure (116±7 vs. baseline: 120±7 mmHg, p>0.05) (Figs. 6D-I). Injections of Kyn into the pFRG did not evoke AE (Figs. 6D-E; 6H).

Bilateral injections of kyn into the pFRG did not change the increase in Dia_EMG_ frequency (58±6, vs. saline+KCN: 71±5 bpm, p>0.05) induced by hypoxia (Fig. 6F). However, blockade of ionotropic glutamatergic receptors into the pFRG reduced the Dia_EMG_ amplitude (0.400±0.163, vs. saline+KCN: 1.714±0.408 mV, p<0.05) and completely eliminate the AE generated by the Abd_EMG_ (0.000±0.000, vs. saline+KCN: 0.386±0.080 mV, p<0.001) induced by hypoxia (Figs. 6D-H). Kyn within the pFRG did not change the increase in MAP elicited by KCN (142±7, vs. saline+KCN: 146±9 mmHg, p>0.05) (Fig. 6I). In the same magnitude, blockade of ionotropic glutamatergic receptors at pFRG region did not change the increase in Dia_EMG_ frequency (54±6, vs. saline+10% CO_2_: 68±3 bpm, p>0.05), Dia_EMG_ amplitude (0.543±0.285, vs. saline+10% CO_2_: 1±0.234 mV, p>0.05), but completely eliminate the AE generated by the Abd_EMG_ (0.000±0.000, vs. saline+10% CO_2_: 0.271±0.042 mV, p<0.001) induced by hypercapnia (Figs. 6D; 6F-H). Bilateral injections of kyn into the pFRG did not change the increase in MAP elicited by hypercapnia (Fig. 6I).

### Changes in the active expiration elicited by hypoxia after anti-DβH-SAP injections into the C1 region

The whole set of experiments performed in the present study demonstrated a possible role of C1 cells in the control of expiratory activity mediated by glutamatergic receptors at the level of pFRG. The last experimental protocol was aimed to evaluate the role of catecholaminergic neurons located in the RVLM on the expiratory activity elicited by hypoxia or hypercapnia challenges. We used a toxin conjugated with DβH-SAP that was bilaterally injected into the rostral aspect of ventrolateral medulla in order to produce a depletion of the TH-expressing neurons in the RVLM (Figs. 7A-C). Only rats with the intraparenchymal anti-DβH-SAP injections confined to the RVLM were used. In the control rats, RVLM TH-ir profiles (presumably C1 cells) were found within the ventral aspect of all 6 brainstem levels examined (Fig. 7C). Rats treated with anti-DβH-SAP had a depletion of the C1 neurons located in the RVLM (Figs. 7B-C). In the 6 rats treated with anti-DβH-SAP, counts of TH-ir between −11.60 and −12.80 mm revealed an average depletion of 71±3% (range between 61-80%) (Fig. 7C). At the intermediate and caudal medullary levels (12.80 to −15.20 mm caudal to bregma), the number of TH-ir neurons was unaffected by treatment with anti-DβH-SAP (data not shown). The TH-ir neurons in the dorsal aspect of the medulla (A2 region) or in the ventrolateral pons (A5 region) did not appear to be reduced by the anti-DβH-SAP treatment (data not shown).

**Figure 7).**
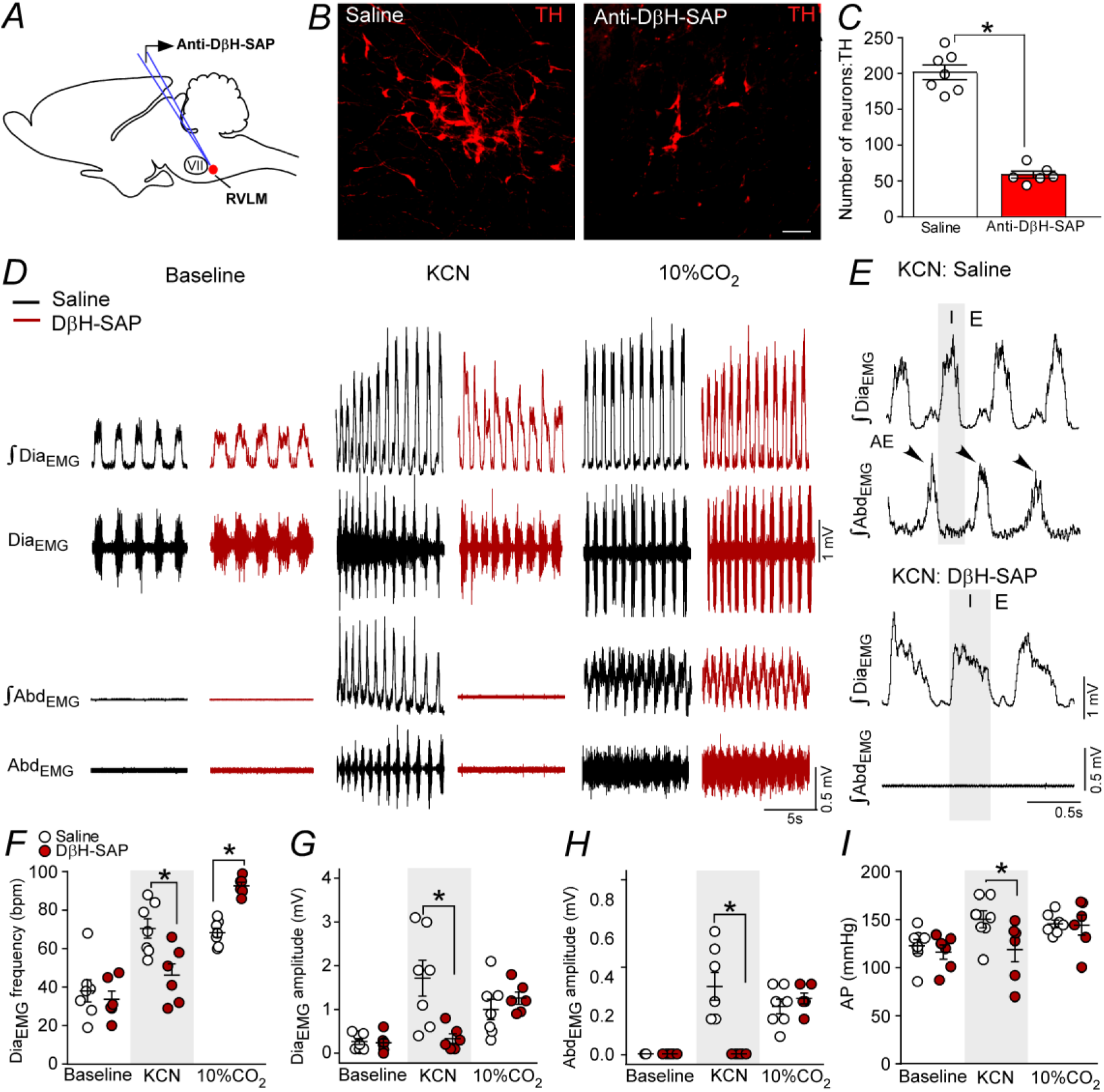
Ablation of the catecholaminergic C1 neurons blunted active expiration induced by hypoxia, but not by hypercapnia. A) Experimental design. B) Photomicrography showing the C1 regions after bilateral injection of saline or anti-DβH-SAP (2.4 ng/100 nL) into the RVLM. C) Number of TH neurons into the RVM of the saline and anti-DβH-SAP-treated groups. D) Traces showing breathing parameters from a representative experiment. E) Expanded traces. Black arrows show active expiration (AE) in the expanded traces. F-I) Individual data for each parameter: F) Dia_EMG_ frequency (bpm), G) Dia_EMG_ amplitude (mV), H) Abd_EMG_ amplitude (mV) and I) AP (mmHg) after bilateral injection of saline (white circles) or anti-DβH-SAP (red circles) into the RVLM. Lines show mean and SEM. *Different from saline; Two-way ANOVA, p<0.05. Abbreviations: RVLM, rostral ventrolateral medulla; Anti-DβH-SAP, immunotoxin anti-dopamine β-hydroxylase-saporin; Dia_EMG_, diaphragm electromyogram; Abd_EMG_, abdominal electromyogram; bpm, breaths per minute; AP, arterial pressure; KCN, potassium cyanide. Scale bar in B = 25 μm.

Selective depletion of catecholaminergic neurons in RVLM did not change breathing parameters at rest (Figs. 7D; 7F-H). As expected, KCN or hypercapnia (10% CO_2_, balanced with O_2_) caused an increase in Dia_EMG_ frequency (KCN: 70±5; CO_2_: 68±2, vs. baseline: 38±6 bpm, p<0.05), Dia_EMG_ amplitude (KCN: 1.714±0.407; CO_2_: 1±0.234 vs. baseline: 0.264±0.067 mV, p<0.05), generate AE in the Abd_EMG_ (KCN: 0.386±0.080; CO_2_: 0.271±0.042, vs. baseline: 0.000±0.000 mV, p<0.05) and increase in MAP (KCN: 146±9, vs. baseline: 120±7 mmHg, p<0.05) (Figs. 7F-I).

Depletion of catecholaminergic neurons of the RVLM blunted the increase in Dia_EMG_ frequency (46±6, vs. saline: 70±5 bpm, p<0.05), Dia_EMG_ amplitude (0.386±0.079, vs. saline: 0.000±0.000 mV, p<0.05) and Abd_EMG_ activity (0.333±0.111, vs. saline: 1.714±0.407 mV, p<0.05) and pressor response (146±9, vs. saline: 118±13 mmHg, p<0.05) elicited by KCN (Figs. 7D-I). On the other hand, depletion of catecholaminergic neurons of the RVLM exacerbates the increase in the Dia_EMG_ frequency (93±2, vs. saline: 68±3 bpm, p<0.05) (Fig. 7F). Anti-DβH-SAP into the RVLM did not affect Dia_EMG_ amplitude (1.258±0.139, vs. saline: 1±0.234 mV, p>0.05) and Abd_EMG_ activity (0.316±0.031, vs. saline: 0.271±0.042 mV, P>0.05) nor the pressor response (141±10, vs. saline: 141±4 bpm, P>0.05) elicited by hypercapnia (Figs. 7D-I).

Injections of the toxin anti-DβH-SAP located outside the C1 region often reached rostral aspects of the ventral lateral medulla, i.e a region that harbors the chemically-coded Phox2b^+^/TH^−^ neurons of the retrotrapezoid nucleus (N = 5), and facial motor nucleus (N = 2). The misplaced injections did not reach caudal region of the ventrolateral medulla, i.e catecholaminergic A1 region. Bilateral injections of the toxin anti-DβH-SAP outside the C1 region produce no significant changes in the breathing parameters under normoxia, cytotoxic hypoxia or hypercapnia condition (data not shown).

## Discussion

Summarizing the results presented in this manuscript, we found a connection between catecholaminergic C1 neurons and pFRG, which is essential for the generation of active expiration under hypoxic condition. In addition, it seems that C1 neurons release glutamate at the level of pFRG in order to trigger expiratory activity. Previous studies (Marina *et al.*, 2011; Abbott *et al.*, 2013*b*; Burke *et al.*, 2014; Malheiros-Lima *et al.*, 2017; Menuet *et al.*, 2017; Malheiros-Lima *et al.*, 2018*b*) and the present findings suggest that, in the intact adult brain, the C1 neurons recruit inspiratory and expiratory muscles in an orderly sequence that depends on the degree to which they are activated by hypoxia or by other mechanisms.

### Catecholaminergic C1 neurons project to pFRG

We found a significant number of fibers and varicosities within the pFRG region. Our transductions were restricted to C1 region, indicating that GFP fibers and varicosities located in the pFRG belong to C1 neurons residing in the rostral aspect of the ventrolateral medulla. The strongest evidence that C1 neurons in fact project to pFRG is the presence of GFP and VGLUT2 double-labeling in this region. TH and CTb double-labeling in the C1 region, observed in rats that received retrograde tracer injection (CTb) into pFRG, reinforce our hypothesis. Therefore, our data show that C1 neurons project to pFRG, and suggest that they could modulate the pFRG activity by releasing both glutamate and catecholamines.

Catecholaminergic neurons from A2/C2 project to parafacial region of the brainstem (Viemari & Ramirez, 2006; Doi & Ramirez, 2010). Therefore, the fibers and terminals only labeled with TH in the pFRG could derivate from other catecholaminergic regions.

### Hypoxia triggering active expiration depends on C1 cells

Active expiration is well defined as the recruitment of expiratory muscles for breathing which occurs during high metabolic demand such as exercise, hypercapnia or hypoxia (Iscoe, 1998). Opinions vary regarding the role of parafacial region (ventral x lateral sub-regions) in this process (Janczewski & Feldman, 2006; Marina *et al.*, 2010; Pagliardini *et al.*, 2011; Abbott *et al.*, 2014; Britto & Moraes, 2017; Zoccal *et al.*, 2018). In addition, anatomical origin, mechanism of generation and modulation of the breathing rhythm has been, and continues to be, intensely investigated (Del Negro *et al.*, 2018). Our results add the information that the recruitment of active expiration under hypoxia depends on the integrity of C1 cells.

As described before, active expiration may also be generated in hypoxic conditions (Abdala *et al.*, 2009; Malheiros-Lima *et al.*, 2017). We propose the existence of a catecholaminergic mechanism for the generation of active expiration since selective ablation of catecholaminergic C1 neurons in the rostral ventrolateral medulla attenuated the late-expiratory flow and the late-E peak of Abd_EMG_ activity observed during hypoxia in non-anesthetized (Abbott *et al.*, 2014; Malheiros-Lima *et al.*, 2017) and anesthetized rats (present study). This effect seems to be exclusive to hypoxia, because ablation of the C1 cells did not affect the generation of active expiration elicited by hypercapnia. These results suggest that recruitment of Abd muscles by chemosensory drives could be mediated through different neuromodulatory systems (Leirão *et al.*, 2018; O’Halloran, 2018).

The control of expiratory activity by C1 neurons is consistent with their function of increasing lung ventilation according to the intensity of the stimulation, hypercapnic or hypoxic stimulus (Guyenet & Bayliss, 2015; Pisanski & Pagliardini, 2019). The former and present data do not clarify whether inspiratory and expiratory activities are regulated by the same C1 neurons, however active expiration triggered by hypoxia depends on the integrity of C1 neurons located in the rostral aspect of the ventrolateral medulla. The main question that needs to be solved is how these neurons are involved in breathing regulation, specially under high metabolic demand such as hypoxia. A direct connection between the C1 cells and elements of the ventral respiratory group is already described (Lipski *et al.*, 1995; Agassandian *et al.*, 2012; Abbott *et al.*, 2013*a*; Burke *et al.*, 2014; Stornetta *et al.*, 2016; Menuet *et al.*, 2017; Malheiros-Lima *et al.*, 2018*a*). C1 neurons are activated by hypoxia and then activate pFRG neurons through a direct glutamatergic mechanism in order to trigger active expiration. The hypoxic activation of C1 neurons is relayed by a direct input from the commissural nucleus of the solitary tract, which receives carotid body afferents (Chitravanshi & Sapru, 1995; Aicher *et al.*, 1996; Koshiya & Guyenet, 1996; Moreira *et al.*, 2006). The activation of C1 neurons by hypoxia can activate breathing through connections with the respiratory column, including the respiratory rhythm-generating neurons located in the pre-Bötzinger complex (Pilowsky *et al.*, 1990; Kang *et al.*, 2017; Malheiros-Lima *et al.*, 2018*a*), inspiratory premotor neurons in the rVRG (Card *et al.*, 2006; Guyenet *et al.*, 2013; Burke *et al.*, 2014) and the pFRG region (present results). We should also consider other brainstem regions involved in breathing regulation such as the parabrachial/Kolliker-Fuse complex (Song & Poon, 2009; Damasceno *et al.*, 2014; Silva *et al.*, 2016*a*), and NTS (Gozal *et al.*, 1999; Song *et al.*, 2011).

As expected, we did not observe significant changes in MAP after pharmacological manipulation of the pFRG region which is consistent with the fact that the lateral aspect of the parafacial region have been proposed to be critical for expiratory rhythm generation (Janczewski & Feldman, 2006; Pagliardini *et al.*, 2011; Huckstepp *et al.*, 2015, 2016).

Our work, taken together with previous work from others, indicates that catecholaminergic neurons in the C1 region use glutamate primarily, if not exclusively, as a transmitter to influence downstream autonomic and/or respiratory neurons.

### Histology and experimental limitations

As shown previously (Malheiros-Lima *et al.*, 2018), most neurons that expressed GFP (94%) were catecholaminergic. The transduced catecholaminergic neurons were likely C1 (adrenergic) rather than A1 (noradrenergic) neurons based on: i) anatomical location, ii) expression of VGLUT2 immunoreactive (Stornetta *et al.*, 2002; DePuy *et al.*, 2013; Malheiros-Lima *et al.*, 2018*a*) and iii) catecholaminergic projections replicate the projections of the C1 cells described by others (Card *et al.*, 2006; Menuet *et al.*, 2017; Malheiros-Lima *et al.*, 2018*a*).

It is also important to point out that GFP was expressed by Phox2b-containing neurons consisting of non-catecholaminergic (RTN-chemosenstitive) neurons (Abbott *et al.*, 2009*a*). However, very few transfected neurons were within the medullary surface under the facial motor nucleus where almost every neuron is Phox2b-positive (Takakura *et al.*, 2014).

### Conclusion

In this study, we described that C1 cells are activated by hypoxia and send projections to the expiratory oscillator located into the pFRG region in order to trigger active expiration. Once activated by hypoxia, C1 cells release glutamate to activate pFRG neurons and generate active expiration, which is an emergent respiratory property to maintain physiological homeostasis under high metabolic demand. Therefore, our study provides evidence that these two populations interact in a way that glutamatergic inputs from the C1 to the pFRG are necessary for the activation of the expiratory neurons during hypoxia, which is responsible for the generation of active expiratory pattern. The revealed C1 and pFRG microcircuitry helps us to understand the neural organization of respiratory pattern generator in conditions of hypoxia. Thereby, our findings have potential implications for understanding the development mechanisms for matching respiratory supply and demand during hypoxia in high altitude populations.

## Acknowledgements

This work was supported by the São Paulo Research Foundation (FAPESP; grants: 2016/23281-3 to ACT; 2015/23376-1 to TSM) and the Conselho Nacional de Desenvolvimento Científico e Tecnológico (CNPq; grant: 408647/2018-3 to ACT). FAPESP fellowship (2014/07698-6 and 2017/08696-5 to MRML and 2014/23418-3 to JNS) and CNPq fellowship (301219/2016-8 to ACT and 301904/2015-4 to TSM). This study was also financed in part by the Coordenação de Aperfeiçoamento de Pessoal de Nível Superior - Brasil (CAPES) - Finnancial Code 001.

## Authors contributions

MRML, JNS, ACT and TSM designed the research; MRML, JNS and FCS performed the research; MRML and JNS analyzed the data; MRML, ACT and TSM wrote the paper.

## References

Abbott SBG, Coates MB, Stornetta RL & Guyenet PG (2013a). Optogenetic stimulation of C1 and retrotrapezoid nucleus neurons causes sleep state – dependent cardiorespiratory stimulation and arousal in rats. Hypertension 61, 835–841.

Abbott SBG, Depuy SD, Nguyen T, Coates MB, Stornetta RL & Guyenet PG (2013b). Selective optogenetic activation of rostral ventrolateral medullary catecholaminergic neurons produces cardiorespiratory stimulation in conscious mice. J Neurosci 33, 3164–3177.

Abbott SBG, Holloway BB, Viar KE & Guyenet PG (2014). Vesicular glutamate transporter 2 is required for the respiratory and parasympathetic activation produced by optogenetic stimulation of catecholaminergic neurons in the rostral ventrolateral medulla of mice in vivo. Eur J Neurosci 39, 98–106.

Abbott SBG, Stornetta RL, Fortuna MG, Depuy SD, West GH, Harris TE & Guyenet PG (2009a). Photostimulation of retrotrapezoid nucleus Phox2b-expressing neurons in vivo produces long-lasting activation of breathing in rats. J Neurosci 29, 5806–5819.

Abbott SBG, Stornetta RL, Socolovsky CS, West GH & Guyenet PG (2009b). Photostimulation of channelrhodopsin-2 expressing ventrolateral medullary neurons increases sympathetic nerve activity and blood pressure in rats. J Physiol 587, 5613–5631.

Abdala APL, Rybak IA, Smith JC & Paton JFR (2009). Abdominal expiratory activity in the rat brainstem – spinal cord in situ: patterns, origins and implications for respiratory rhythm generation. J Physiol 587, 3539–3559.

Agassandian K, Shan Z, Raizada M, Sved AF & Card JP (2012). C1 catecholamine neurons form local circuit synaptic connections within the rostroventrolateral medulla of rat. Neuroscience 227, 247–259.

Aicher SUEA, Saravay RH, Cravo S, Jeske I, Morrison SF, Reis DJ & Milner ANITA (1996). Monosynaptic projections from the nucleus tractus solitarii to C1 adrenergic neurons in the rostral ventrolateral medulla: Comparison with input from the caudal ventrolateral medulla. J Comp Neurol 373, 62–75.

Britto AA De & Moraes DJA (2017). Non-chemosensitive parafacial neurons simultaneously regulate active expiration and airway patency under hypercapnia in rats. J Physiol 595, 2043–2064.

Burke PGR, Abbott SBG, Coates MB, Viar KE, Stornetta RL & Guyenet PG (2014). Optogenetic stimulation of adrenergic C1 neurons causes sleep state – dependent cardiorespiratory stimulation and arousal with sighs in rats. Am J Respir Crit Care Med 190, 1301–1310.

Card JP, Sved JC, Craig B, Raizada M, Vazquez J & Sved AF (2006). Efferent projections of rat rostroventrolateral medulla C1 catecholamine neurons: implications for the central control of cardiovascular regulation. J Comp Neurol 499, 840–859.

Chitravanshi V & Sapru HN (1995). Chemoreceptor-sensitive subnucleus of nucleus neurons in commissural tractus solitarius of the rat. Am J Physiol - Regul Integr Comp Physiol 268, R851–858.

Damasceno RS, Takakura AC & Moreira TS (2014). Regulation of the chemosensory control of breathing by Kölliker-Fuse neurons. Am J Physiol - Regul Integr Comp Physiol 307, R57–R67.

Dampney RAL (1994). Functional organization of central pathways regulating the cardiovascular system. Physiol Rev 74, 323–364.

DePuy SD, Stornetta RL, Bochorishvili G, Deisseroth K, Witten I, Coates M & Guyenet PG (2013). Glutamatergic neurotransmission between the C1 neurons and the parasympathetic preganglionic neurons of the dorsal motor nucleus of the vagus. J Neurosci 33, 1486–1497.

Doi A & Ramirez J (2010). State-dependent interactions between excitatory neuromodulators in the neuronal control of breathing. J Neurosci 30, 8251–8262.

Gozal D, Xue Y & Simakajornboon N (1999). Hypoxia induces c-Fos protein expression in NMDA but not AMPA glutamate receptor labeled neurons within the nucleus tractus solitarii of the conscious rat. Neurosci Lett 262, 93–96.

Guyenet P, Stornetta R, Bochorishvili G, DePuy S, Burke P & Abbott S (2013). C1 neurons: The body’s EMTs. Am J Physiol - Regul Integr Comp Physiol 305, R187–R204.

Guyenet PG (2000). Neural structures that mediate sympathoexcitation during hypoxia. Respir Physiol 121, 147–162.

Guyenet PG (2006). The sympathetic control of blood pressure. Nat Rev Neurosci 7, 335–346.

Guyenet PG (2014). Regulation of breathing and autonomic outflows by chemoreceptors. Compr Physiol 4, 1511–1562.

Guyenet PG & Bayliss DA (2015). Neural Control of Breathing and CO2 Homeostasis. Neuron 87, 946–961.

Guyenet PG, Mulkey DK, Stornetta RL & Bayliss DA (2005). Regulation of ventral surface chemoreceptors by the central respiratory pattern generator. J Neurosci 25, 8938–8947.

Huckstepp RTR, Cardoza KP, Henderson LE & Feldman JL (2018). Distinct parafacial regions in control of breathing in adult rats. PLoS One 13:e020148, 1–24.

Huckstepp RTR, Cardoza KP, Henderson LE & Feldman XL (2015). Role of parafacial nuclei in control of breathing in adult rats. J Neurosci 35, 1052–1067.

Huckstepp RTR, Henderson LE, Cardoza KP & Feldman JL (2016). Interactions between respiratory oscillators in adult rats. Elife 5:e14203, 1–22.

Hwang D, Carlezon WA, Isacson OLE & Kim K (2001). A high-efficiency synthetic promoter that drives transgene expression selectively in noradrenergic neurons. Hum Gene Ther 12, 1731–1740.

Iscoe S (1998). Control of abdominal muscles. Prog Neurobiol 56, 433–506.

Janczewski WA & Feldman JL (2006). Distinct rhythm generators for inspiration and expiration in the juvenile rat. J Physiol 570, 407–420.

Kang J, Liang W, Lam C, Huang X, Yang S, Wong-Riley M, Fung M & Liu Y (2017). Catecholaminergic neurons in synaptic connections with pre-Bötzinger complex neurons in the rostral ventrolateral medulla in normoxic and daily acute intermittent hypoxic rats. Exp Neurol 287, 165–175.

King TL, Kline DD, Ruyle BC, Heesch CM & Hasser EM (2013). Acute systemic hypoxia activates hypothalamic paraventricular nucleus-projecting catecholaminergic neurons in the caudal ventrolateral medulla. Am J Physiol - Regul Integr Comp Physiol 305, R1112–R1123.

Koshiya N & Guyenet PG (1996). Tonic sympathetic chemoreflex after blockade of respiratory rhythmogenesis in the rat. J Physiol 491, 859–869.

Leirão I, Silva Jr C, Gargaglioni LH & Silva GSF (2018). Hypercapnia-induced active expiration increases in sleep and enhances ventilation in unanaesthetized rats. J Physiol 596, 3271–3283.

Li Y-W & Guyenet PG (1996). Angiotensin II Decreases a Resting K+ Conductance in Rat Bulbospinal Neurons of the C1 Area. Circ Res 78, 274–282.

Lipski J, Kanjhan R, Kruszewska B & Smith M (1995). Barosensitive neurons in the rostral ventrolateral medulla of the rat in vivo: morphological properties and relationship to C1 adrenergic neurons. Neuroscience 69, 601–618.

Malheiros-Lima MR, Takakura AC & Moreira TS (2017). Depletion of rostral ventrolateral medullary catecholaminergic neurons impairs the hypoxic ventilatory response in conscious rats. Neuroscience 351, 1–14.

Malheiros-Lima MR, Totola LT, Lana MVG, Strauss BE, Takakura AC & Moreira TS (2018a). Breathing responses produced by optogenetic stimulation of adrenergic C1 neurons are dependent on the connection with preBötzinger complex in rats. Pflugers Arch Eur J Physiol 470, 1659–1672.

Malheiros-Lima MR, Totola LT, Takakura AC & Moreira TS (2018b). Impaired chemosensory control of breathing after depletion of bulbospinal catecholaminergic neurons in rats. Pflugers Arch Eur J Physiol 470, 277–293.

Marina N, Abdala AP, Trapp S, Li A, Nattie EE, Hewinson J, Smith JC, Paton JFR & Gourine A V (2010). Essential role of Phox2b-expressing ventrolateral brainstem neurons in the chemosensory control of inspiration and expiration. J Neurosci 30, 12466–12473.

Marina N, Abdala APL, Korsak A, Simms AE, Allen AM, Paton JFR & Gourine A V (2011). Control of sympathetic vasomotor tone by catecholaminergic C1 neurones of the rostral ventrolateral medulla oblongata. Cardiovasc Res 91, 703–710.

McAllen RM & Dampney RAL (1990). Vasomotor neurons in the rostral ventrolateral medulla are organized topographically with respect to type of vascular bed but not body region. Neurosci Lett 110, 91–96.

Menuet C, Le S, Dempsey B, Connelly A, Kamar J, Jancovski N, Bassi J, Walters K, Simms A, Hammond A, Fong A, Goodchild A, Mcmullan S & Allen A (2017). Excessive respiratory modulation of blood pressure triggers hypertension. Cell Metab 25, 739–748.

Moreira TS, Takakura AC, Colombari E & Guyenet PG (2006). Central chemoreceptors and sympathetic vasomotor outflow. J Physioll 577, 369–386.

Del Negro CA, Funk GD, Feldman JL & Angeles L (2018). Breathing matters. Nat Rev Neurosci 19, 351–367.

O’Halloran K (2018). Sleep awakens active expiration. J Physiol 596, 2947–2948.

Pagliardini S, Janczewski WA, Tan W, Dickson CT, Deisseroth K & Feldman JL (2011). Active expiration induced by excitation of ventral medulla in adult anesthetized rats. J Neurosci 31, 2895–2905.

Pattyn A, Morin X, Cremer H, Goridis C & Brunet J (1997). Expression and interactions of the two closely related homeobox genes Phox2a and Phox2b during neurogenesis. Development 124, 4065–4075.

Pilowsky PM, Jiang C & Lipski J (1990). An intracellular study of respiratory neurons in the rostral ventrolateral medulla of the rat and their relationship to catecholamine-containing neurons. J Comp Neurol 301, 604–617.

Pisanski A & Pagliardini S (2019). The parafacial respiratory group and the control of active expiration. Respir Physiol Neurobiol 265, 153–160.

Prabhakar NR & Semenza GL (2015). Oxygen Sensing and Homeostasis. Physiology 30, 340–348.

Ritter S, Llewellyn-Smith I & Dinh TT (1998). Subgroups of hindbrain catecholamine neurons are selectively activated by 2-deoxy-D-glucose induced metabolic challenge. Brain Res 805, 41–54.

Ross CA, Armstrong DM, Ruggiero D, Pickel V, Joh TH & Reis DJ (1981). Adrenaline neurons in the rostral ventrolateral medulla innervate thoracic spinal cord: a combined immunocytochemical and retrograde transport demonstration. Neurosci Lett 25, 257–262.

Schreihofer A & Guyenet P (1997). Identification of C1 presympathetic neurons in rat rostral ventrolateral medulla by juxtacellular labeling in vivo. J Comp Neurol 387, 524–536.

Silva JN, Lucena E V, Silva TM, Damasceno RS, Takakura AC & Moreira TS (2016a). Inhibition of the pontine Kölliker-Fuse nucleus reduces genioglossal activity elicited by stimulation of the retrotrapezoid chemoreceptor neurons. Neuroscience 328, 9–21.

Silva JN, Oliveira LM, Souza FC, Moreira TS & Takakura AC (2019). Distinct pathways to the parafacial respiratory group to trigger active expiration in adult rats. Am J Physiol Cell Mol Physiol 317, L402–L413.

Silva TM, Takakura AC & Moreira TS (2016b). Acute hypoxia activates hypothalamic paraventricular nucleus-projecting catecholaminergic neurons in the C1 region. Exp Neurol 285, 1–11.

Song G & Poon C-S (2009). Lateral parabrachial nucleus mediates shortening of expiration and increase of inspiratory drive during hypercapnia. Respir Physiol Neurobiol 165, 9–12.

Song G, Xu H, Wang H, MacDonald S & Poon C-S (2011). Hypoxia-excited neurons in NTS send axonal projections to Kölliker-Fuse/parabrachial complex in dorsolateral pons. Neuroscience 175, 145–153.

Stornetta RL, Inglis MA, Viar KE & Guyenet PG (2016). Afferent and efferent connections of C1 cells with spinal cord or hypothalamic projections in mice. Brain Struct Funct 221, 4027–4044.

Stornetta RL, Moreira TS, Takakura AC, Kang BJ, Chang DA, West GH, Mulkey DK, Bayliss DA & Guyenet PG (2006). Expression of Phox2b by brainstem beurons involved in chemosensory integration in the adult rat. J Neurosci 26, 10305–10314.

Stornetta RL, Sevigny CP & Guyenet PG (2002). Vesicular glutamate transporter DNPI/VGLUT2 mRNA is present in C1 and several other groups of brainstem. J Comp Neurol 444, 191–206.

Takakura AC, Barna F, Cruz JC, Colombari E & Moreira TS (2014). Phox2b-expressing retrotrapezoid neurons and the integration of central and peripheral chemosensory control of breathing in conscious rats. Exp Physiol 99, 571–585.

Taxini C, Gargaglioni L, Takakura A & Moreira T (2011). Control of the central chemoreflex by A5 noradrenergic neurons in rats. Neuroscience 199, 177–186.

Viemari J & Ramirez J (2006). Norepinephrine differentially modulates different types of respiratory pacemaker and nonpacemaker neurons generation depends on a heterogeneous pacemaker population. J Neurophysiol 95, 2070–2082.

Zoccal DB, Silva J, Barnett W, Lemes E, Falquetto B, Colombari E, Molkov Y, Moreira T & Takakura A (2018). Interaction between the retrotrapezoid nucleus and the parafacial respiratory group to regulate active expiration and sympathetic activity in rats. Am J Physiol Cell Mol Physiol 315, L891–L909.

